# Reconstruction and Analysis of a *Kluyveromyces marxianus* Genome-scale Metabolic Model

**DOI:** 10.1101/581587

**Authors:** Simonas Marcišauskas, Boyang Ji, Jens Nielsen

## Abstract

**Background:** *Kluyveromyces marxianus* is a thermotolerant yeast with multiple biotechnological potentials for industrial applications, which can metabolize a broad range of carbon sources, including less conventional sugars like lactose, xylose, arabinose and inulin. These phenotypic traits are sustained even up to 45°C, what makes it a relevant candidate for industrial biotechnology applications, such as ethanol production. It is therefore of much interest to get more insight into the metabolism of this yeast. Recent studies suggested, that thermotolerance is achieved by reducing the number of growth-determining proteins or suppressing oxidative phosphorylation. Here we aimed to find related factors contributing to the thermotolerance of *K. marxianus*.

**Results:** Here, we reported the first genome-scale metabolic model of *Kluyveromyces marxianus*, iSM996, using a publicly available *Kluyveromyces lactis* model as template. The model was manually curated and refined to include missing species-specific metabolic capabilities. The iSM996 model includes 1913 reactions, associated with 996 genes and 1531 metabolites. It performed well to predict the carbon source utilization and growth rates under different growth conditions. Moreover, the model was coupled with transcriptomics data and used to perform simulations at various growth temperatures.

**Conclusions:** *K. marxianus* iSM996 represents a well-annotated metabolic model of thermotolerant yeast, which provide new insight into theoretical metabolic profiles at different temperatures of *K. marxianus*. This could accelerate the integrative analysis of multi-omics data, leading to model-driven strain design and improvement.

## Background

The ascomycetous yeast *Kluyveromyces marxianus* is a prospective microbial host for industrial biotechnology. It is known as a fast growing, Crabtree negative organism, capable of utilizing a broad selection of sugars and having a high secretion capacity for proteins [1, 2]. For these reasons, *K. marxianus* has been successfully applied in numerous studies for endogenous enzymes production including inulinase [3], β-galactosidase [4], β-glucosidase [5] and β-xylosidase [6]. Unlike its sister species *Kluyveromyces lactis, K. marxianus* is thermotolerant, growing up to 45-50°C. Since it has been isolated mainly from dairy products for years, it has GRAS (Generally Regarded as Safe) and QPS (Qualified Presumption of Safety) status, therefore making it suitable for applications in the food and pharma industry.

Thermotolerant hosts may positively impact high temperature industrial bioprocesses in several ways. Firstly, thermotolerance is favourable once simultaneous saccharification and fermentation (SSF) is considered as many amylases and cellulases are more efficient at high temperatures [7]. Secondly, the maintenance costs may be reduced due to lower cost for bioreactor cooling and sterilization of cultivation media [7, 8]. The media sterility is less important at high temperatures due to the decreased competition between the species for available medium resources. This is particularly beneficial for complex media like whey and slurry. At high temperatures, thermotolerant yeasts are forced to reorganize their profiles at the transcriptional and proteomic levels [9]. For instance, protein folding, phosphorylation and cell wall organisation processes were upregulated in YPD medium under high temperature, thereby suppressing carbohydrate, amino acid and lipid metabolism [10, 11]. The cell likely benefits from the medium richness, while depleting biosynthetic pathways for several YPD compounds. However, not much is known about *K. marxianus* intracellular metabolite levels at high temperatures. Such knowledge may be useful for minimal media design and for identifying high temperature metabolic capabilities, in particular for evaluation of the production potential of various compounds.

Genome-scale metabolic models (GEMs) are valuable systems biology tools as they combine genome annotation, cultivation and other experimental data for a given organism into a whole-cell metabolic network. Complementary to general omics analysis, GEMs give an ability to integrate omics data into metabolic networks and predict condition specific metabolic capabilities. GEMs have been successfully used to design strains and evaluate the cell capabilities upon different conditions [12, 13].

To evaluate the metabolic capabilities of *K. marxianus*, we reconstructed iSM996, the first genome-scale metabolic model (GEM) for this species. This model was used to predict the growth on various carbon sources and we found good consistency between model predictions and experimental data reported in the literature. We used transcriptomics data to obtain context specific models at different chemostat conditions, i.e. 30°C YPD shaking, 30°C YPD non-shaking and 45°C YPD shaking. The models revealed the main metabolic bottlenecks associated with growth at high temperatures. To our knowledge, this approach is one of the first attempts to use GEMs to evaluate metabolic profiles at different temperatures.

## Methods

### Model reconstruction

The genome-scale metabolic network reconstruction for *K. marxianus* DMKU3-1042 was based on published genome annotation (NCBI accession PRJDA65233) and other databases including KEGG [14], MetaCyc [15], TransportDB [16] and BRENDA [17]. Two draft models were generated using the RAVEN Toolbox [18]. The first model contained homologous reactions from iOD907, the GEM for the sister species *Kluyveromyces lactis* [19]. This was accomplished with RAVEN function *getModelFromHomology*, which utilises BLASTP [20] for the bi-directional homology search. The following homology criteria were used: e-value 1E-30; identity 40%; alignment length of 200. Additional check to identify homologs was considered for proteins shorter than 250 amino acids, since for them it was less likely to satisfy e-value and alignment length criteria. Spontaneous iOD907 reactions without gene associations were also checked to be added to this draft model. The second draft model was generated from KEGG using RAVEN function *getKEGGModelForOrganism*. This function uses HMMER (http://hmmer.org/) in homology search, where the query proteome is queried against KEGG Orthology (KO) specific hidden Markov model (HMM) sets. The default e-value threshold was used, i.e. 1E-50. It was then compartmentalised with RAVEN function *predictLocalization*, which uses WoLF PSORT [21] protein scores as input. Manual curation was then performed for metabolite names and reactions reversibility, directionality, when using iOD907, Yeast 7.6 [22] and HMR2 [23] as reference. Thereafter, the unique reactions from the second draft model were then manually added to the first draft model. All the compartmentalisation discrepancies were fixed according to the latter model.

### Biomass composition

The biomass composition was adapted from iOD907 and adjusted with *K. marxianus* bibliographic data wherever possible. The cell dry weight composition (g/gDW) for protein, carbohydrate, lipid, RNA and DNA content was incorporated from cultivation study [24]. The stoichiometric coefficients for biomass compounds were then scaled-up to constitute up to 1 gram of biomass in biomass equation. The experimental data from the cell wall study [25] was used to estimate the composition of the cell wall carbohydrates, namely glucan, mannan and chitin. Knowing the total carbohydrates mass in 1 g of biomass, we calculated the remaining mass for other carbohydrates (trehalose and amylose). The molar ratio between trehalose and amylose was considered the same as in iOD907. Nucleotide, deoxynucleotide and amino acid composition was calculated from genome, transcriptome (including tRNA and rRNA) and proteome according to the suggested protocol [12]. The detail biomass composition is shown in Additional file 3. The energy parameters for growth associated maintenance (GAM) and the non-growth associated maintenance (NGAM) and P/O ratio was retained the same as in iOD907, since related information was lacking for *K. marxianus*.

### Manual curation

The semi-automatic gap filling was performed to ensure that the draft model could produce all biomass components in Verduyn medium [26]. This medium was also considered as the minimal medium in this study. The main gap-filling reaction sources were iOD907, Yeast 7.6 models and KEGG database. Such pre-processed draft model was then greatly improved with *K. marxianus* literature knowledge. For instance, we added the missing uptake pathways for several carbon [27] sources like inulin, L-arabinose or D-mannitol. The draft model was then thoroughly curated for gene associations, EC numbers, metabolite names, reactions elemental/charge balance. Here we considered Yeast 7.6 [22], BRENDA [17] and MetaCyc [15], TransportDB [16] as reference. The additional information for every reaction is included in Additional file 2 (the list of reactions, followed by the column which state from in which step it was added).

### Model validation

The model was validated using constraint-based flux balance analysis (FBA) upon minimal medium. An ability to predict growth upon various carbon and nitrogen sources was tested and compared with the literature knowledge. During testing of different nitrogen sources D-glucose was chosen as a carbon source. The iSM996 was also checked for the growth rate accuracy in minimal media including various uptake rates for carbon sources. In such simulations, the lower and upper bounds for relevant substrates were set to the corresponding experimental uptake rate values.

### Transcriptomic data integration

The quantitative TSS-seq data was used to deactivate reactions and obtain condition specific models. Three conditions were considered: 30°C YPD shaking (30D), 30°C YPD non-shaking (30DS) and 45°C YPD shaking (45D). TSS-seq dataset also contained the data for D-xylose medium (YPX) in 30°C (YPX). However, it was decided not to include this condition in the analysis, since KmXKS1 gene, responsible for D-xylulose phosphorylation, was inactive therefore preventing D-xylose assimilation. However, all four conditions were utilised to identify the genes which were turned off in at least one condition. Thereafter, three condition specific models were obtained, each having inactivated reactions once their associated genes were inactive.

## Results and Discussion

### Model reconstruction

The model reconstruction of *K. marxianus* included four steps: i) obtain and curate the draft reconstruction based on template model iOD907; ii) fetch and curate the second draft model from KEGG; iii) merge both draft models; iv) perform the thorough manual curation. The first draft model after curation had 875 genes including 20 manually identified homologs and 29 non-gene-associated spontaneous reactions. Meanwhile, the compartmentalised and curated second draft model had 524 genes. These two models were merged into a new draft model with 987 genes, 1934 reactions and 1678 metabolites. We then added exchange, biomass and ATP maintenance reactions from iOD907 and performed growth simulation in a minimal medium. As the model could predict growth, we used the draft model as a basis for further model modifications.

The manual curation started by fixing two inconsistencies inferred from the template GEM. Firstly, in iOD907 all the cytosolic metabolites could be transported to extracellular space and had exchange reactions. Since the cell ability to import or secrete all cytosolic metabolites seems unspecific, we commenced the manual check by using Yeast 7.6 as reference. Overall, the number of transportable cytosolic metabolites was decreased from 313 to 182 in iSM996. Another iOD907 problem was that the cell could import or secrete several fatty acid-acyl carrier protein (ACP) complexes even though the cell membrane should be impermeable for such large complexes. These complexes were therefore replaced by the corresponding free fatty acids. Correspondingly, the biosynthetic pathways were added for the known products like 2-phenylethanol, phenethyl acetate or ethyl acetate.

The iSM996 model (**Figure 1a**) features 996 genes, 1913 reactions and 1531 metabolites thereby including *K. marxianus* specific 110 genes, 455 reactions and 298 metabolites. The most represented metabolic pathways are related to transport, exchange reactions and amino acid, lipid, carbohydrate metabolism (**Figure 1b**). As shown in **Table 1**, iSM996 has higher genome coverage than iOD907, however, it has lower number of unique (non-S. *cerevisiae*) genes. Regarding the reactions content, iSM996 contains more reactions occurring in cytosol and less in extracellular space.

**Figure 1.**
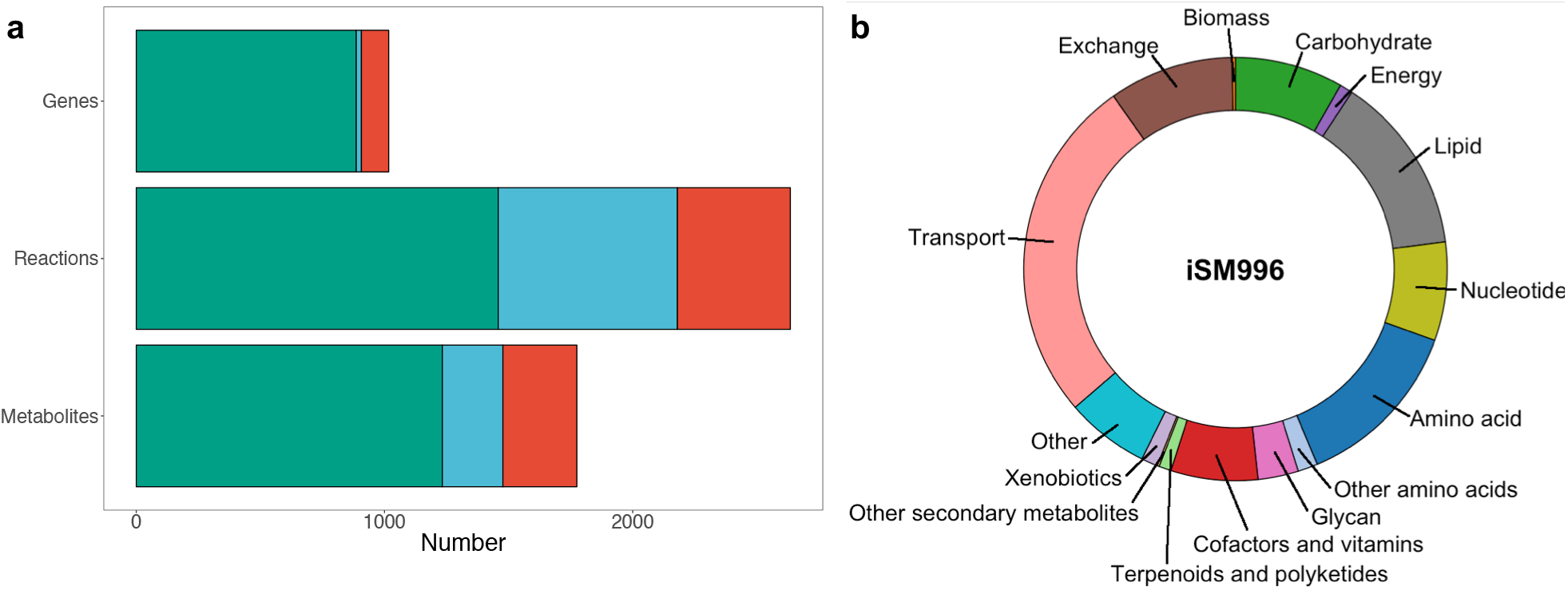
Overview of iSM996. (a) Comparison of genes, reactions and metabolites present in iSM996 and template model (iOD907). Green colour indicates overlapping entities, blue – specific to iOD907, red – specific to iSM996. (b) Distribution of reactions in each metabolic part.

**Table 1.** Comparison between *Kluyveromyces lactis* GEM iOD907 and *Kluyveromyces marxianus* GEM iSM996.

|  | iOD907 | iSM996 |
| --- | --- | --- |
| Genes | 907 (17.8%) | 996 (20.1%) |
| *S. cerevisiae* homologues | 691 | 916 |
| Unique | 216 | 80 |
| Reactions | 2180 | 1913 |
| Extracellular | 938 | 507 |
| Cytosol | 853 | 974 |
| Mitochondria | 359 | 390 |
| Endoplasmic Reticulum | 30 | 42 |
| Metabolites | 1477 | 1531 |
| Extracellular | 313 | 191 |
| Cytosol | 822 | 907 |
| Mitochondria | 296 | 359 |
| Endoplasmic Reticulum | 46 | 74 |

### Model validation

We validated the iSM996 model by checking the growth in known carbon and nitrogen sources in minimal media (**Figure 2a**). The growth could be predicted in numerous monosaccharides (glucose, galactose, D-xylose), disaccharides (sucrose, lactose, cellobiose) and polysaccharides (inulin). The iSM996 could also predict growth in amino acid-free minimal medium, meaning that the cell is capable to *de novo* synthesize all the required amino acids. While *in silico* simulations showed growth upon all nitrogen sources, no growth could be predicted for L-lysine and cadaverine as sole carbon sources. The most likely scenario how L-lysine is catabolised and then directed to the central carbon metabolism is the 6-step linear pathway (MetaCyc LYSDEGII-PWY) occurring in *Saccharomyces cerevisiae* and providing glutarate as the final product. One remaining step would then be to convert glutarate into glutaryl-CoA, which may be further modified by known fungal enzymes. Cadaverine is the product of L-lysine decarboxylation, which does not occur in LYSDEGII-PWY. Since it is unknown how cadaverine is further catabolised in fungi, one may speculate that cadaverine is converted back to L-lysine with carbon fixation and then processed in the same hypothetical way as L-lysine. Due to a high uncertainty regarding these hypotheses, the decision was made not to include these reactions to iSM996.

**Figure 2.**
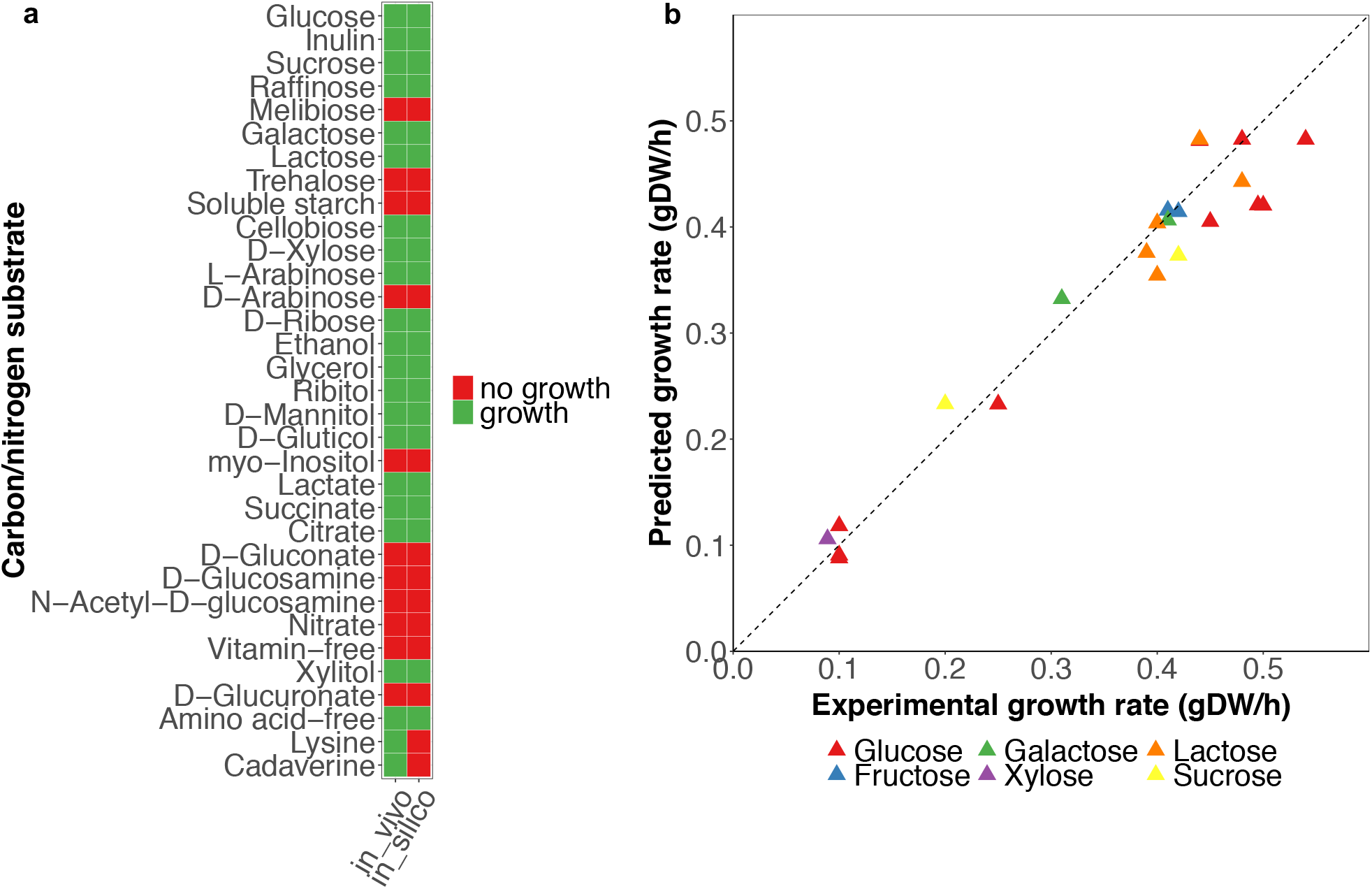
Validation results for iSM996 in minimal medium. (a) Comparison of in silico growth for various carbon and nitrogen sources with literature data. Upon simulations for nitrogen sources it was assumed that glucose is a carbon source. (b) Comparison of *in silico* growth rate and experimental growth rate for various carbon sources in minimal medium. The squared value of Pearson correlation coefficient between experimental and predicted growth values was 0.9445.

The iSM996 was also checked for the growth rate prediction accuracy upon various carbon sources and their uptake rates. The results (**Figure 2b**) indicate the strong correlation between experimental and *in silico* growth rates. As expected, the difference between *in vivo* and *in silico* growth rates was smaller when growth rate did not exceed 0.3 h^-1^. Whereas NGAM value has a minimal impact to predicted growth rates, one may hypothesize that in reality the cell “tweaks” the stoichiometric coefficients in the biomass equation including GAM. It is therefore not possible to re-use the same biomass composition in a wide range of the growth rates. While biomass reactions in GEMs are usually most relevant for growth rates lower than 0.4 h^-1^, more specific experimental data for the biomass composition, GAM, and NGAM are needed for higher growth rates, if one aims to perform simulations at high growth rates.

### *In silico* evaluation of *K. marxianus* metabolic capacities in stress conditions

The iSM996 model represents the whole cell metabolic capabilities assuming that all genes are active. We next used iSM996 to predict the cell metabolic stress response upon low oxygen availability and high temperatures in rich medium. This was done by integrating TSS-seq data to GEM and obtaining context-specific models in YPD medium for the following conditions: 30°C shaking (30D), 30°C static (30DS) and 45°C shaking (45D).

The TSS-seq data was used to turn off gene-associated reactions without transcriptional evidence while leaving constraints for other reactions intact. These models were also used to predict the maximal production capacity for the main biomass components (**Figure 3**). We calculated the excessive production above the metabolic demands for achieving 90% of the maximal growth rate. TSS-seq data hinted towards a noticeably tighter metabolic regulation in 45D (80 inactivated genes) compared with 30D and 30DS (24 and 15 inactivated genes respectively). The initial FBA simulations revealed the riboflavin auxotrophy in 30DS and 45D conditions, and ferroheme auxotrophy 45D condition. To enable metabolic profile comparison between the conditions it was assumed that these metabolites are available and can be transported from the medium into the cell. Upon such consideration, the growth was predicted in all three condition-specific models. Reduction cost analysis showed that the growth was limited by L-cysteine availability and this represented a limitation at all three conditions. The high temperature condition therefore featured the most zero-flux reactions (638) while 30D and 30DS correspondingly had 544 and 541 such reactions.

**Figure 3.**
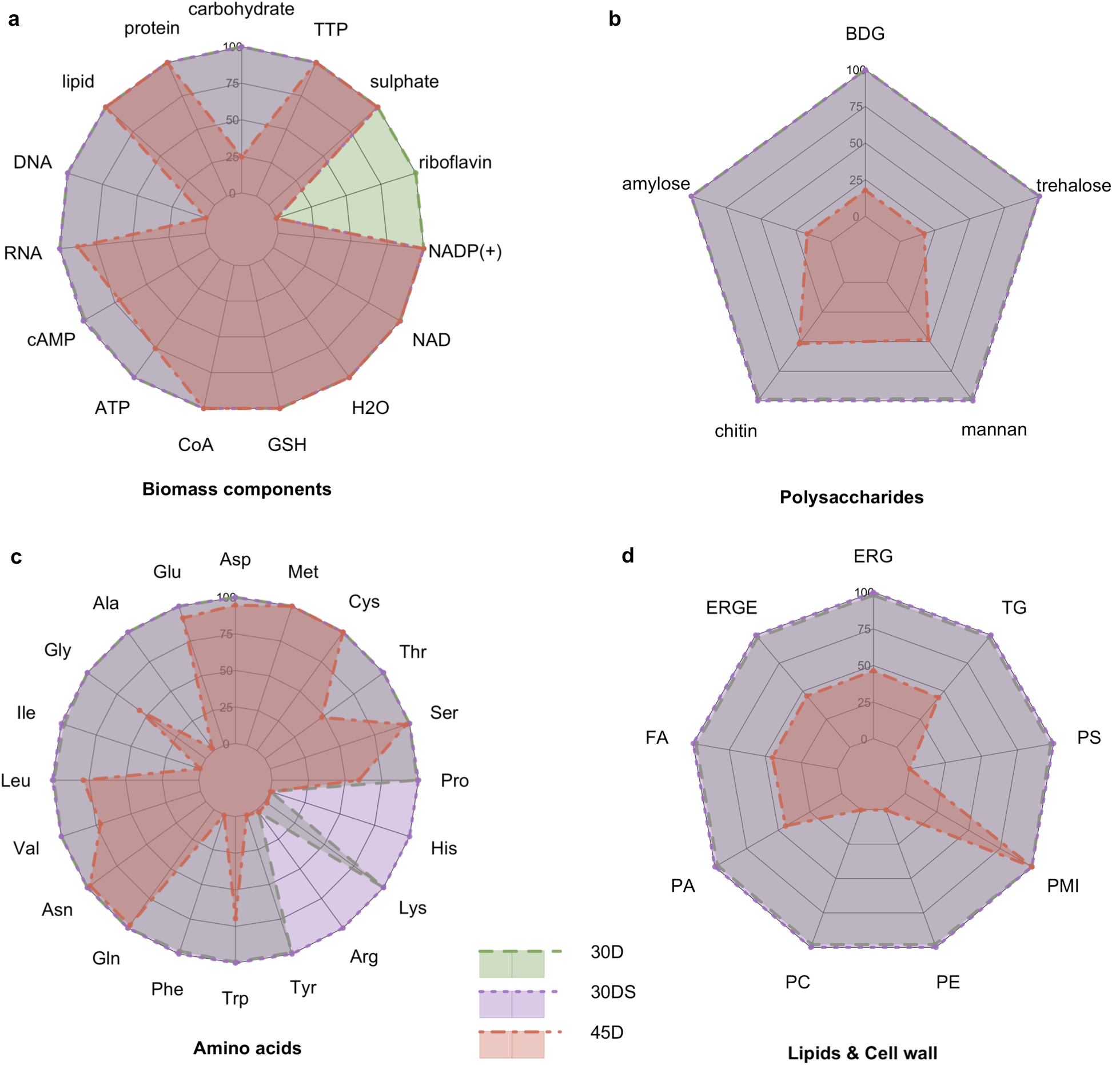
A radar chart showing the predicted potential for biomass precursors excessive production in 30D, 30DS and 45D conditions. As the magnitude is different for each metabolite, the relative production values are shown, where 100% indicates the largest production capacity between conditions. The data for the 30D condition is shown as the green polygon bordered with the dashed border while the corresponding data for the 30DS condition is in purple (dotted border) and the data for the 45D condition is in red (dot dash border) color. (a) Abbreviations: cAMP (3’,5’-cyclic AMP), CoA (coenzyme A), GSH (reduced glutathione), TTP (deoxythymidine 5’-triphosphate). (b) Abbreviations: BDG ((1->3)-beta-D-glucan). (c) Abbreviations (by side chain class): a) acid: Asp (L-aspartate), Glu (L-glutamate); b) aliphatic: Ala (L-alanine), Gly (glycine), Ile (L-isoleucine), Leu (L-leucine), Val (L-valine); c) amide: Asn (L-asparagine), Gln (L-glutamine); d) aromatic: Phe (L-phenylalanine), Trp (L-tryptophan), Tyr (L-tyrosine); e) basic: Arg (L-arginine), Lys (L-lysine); f) basic aromatic: His (L-histidine); g) Pro (L-proline); hydroxyl-containing: Ser (L-serine), Thr (L-threonine); h) sulphur containing: Cys (L-cysteine), Met (L-methionine). (d) Abbreviations: ergosterol (ERG), ergosterol ester (ERGE), FA (fatty acid), PA (phosphatidate), PC (phosphatidylcholine), PE (phosphatidylethanolamine), PMI (1-phosphatidyl-1D-myo-inositol), PS (phosphatidyl-L-serine), TG (triglyceride). The corresponding radar charts for precursor metabolites nucleotides are included in Figure S1.

The simulations indicate that across the conditions, *K. marxianus* retains the ability to allocate amino acids necessary for roughly the same protein mass. Amino acid auxotrophies to L-arginine and L-histidine were identified in 30D and 45D. High temperature-specific auxotrophies included the previously reported L-lysine, L-isoleucine [28] and previously undiscovered auxotrophies to L-alanine, L-phenylalanine and L-tyrosine. Having depleted biosynthetic pathways for seven amino acids at high temperatures allows the cells to conserve precursor metabolites required for these pathways and likely contributes to an increased flux through Embden–Meyerhof–Parnas (EMP) pathway and the TCA cycle due to the increased carbon availability. However, the decreased theoretical production pool sizes in 45D for L-alanine, L-isoleucine, L-phenylalanine, L-tyrosine and L-arginine may reduce the availability for biosynthesis of some proteins enriched by these amino acids.

The auxotrophy to riboflavin, found in 30DS and 45D conditions, has other potential benefits in addition to just conserving building blocks. Two condition-specific strategies are considered to inactivate *de novo* riboflavin synthesis. As shown in **Figure 4**, the main riboflavin precursors GTP and D-ribulose 5-phosphate are processed in their corresponding linear pathways until their final products are used to produce 6,7-dimethyl-8-(1-D-ribityl)lumazine, the immediate precursor for riboflavin. Upon 30DS condition, the expression of *KmRIB3*, which catalyses the final reactions of GTP and D-ribulose 5-phosphate linear pathways (r_0939 and r_0940 correspondingly) is inactivated. However, the interconversion ability is possibly retained between riboflavin and the downstream metabolites FMN and FAD. At the 45D condition, the expression of *KmRIB4* and *KmFMN1* genes were inactivated. The former gene is responsible for 6,7-dimethyl-8-(1-D-ribityl)lumazine production (r_0938), while the latter gene is involved in FMN conversion from riboflavin (r_0937). Thus, under high temperature condition, *K. marxianus* can only interconvert FMN and FAD or hydrolyse it back to riboflavin. These findings suggest that the organism has to find an optimal balance between ATP synthesis (riboflavin) and other important metabolic functions linked with FMN and FAD. While at the 30DS conditions, the cell can freely allocate these three compounds. Under 45D condition, *K. marxianus* features irreversible riboflavin production from FMN and FAD, suggesting that ATP shortage is likely the main limitation for the cell at high temperatures, probably further affected by limited ferroheme availability. The additional supplementation of riboflavin is known to increase the tolerance to high temperatures upon low ATP/ADP ratio [29]. Since the ATP production differences between 30°C and 45°C are only about 20%, one may conclude that an increased ATP demand is related to non-growth associated maintenance processes.

**Figure 4.**
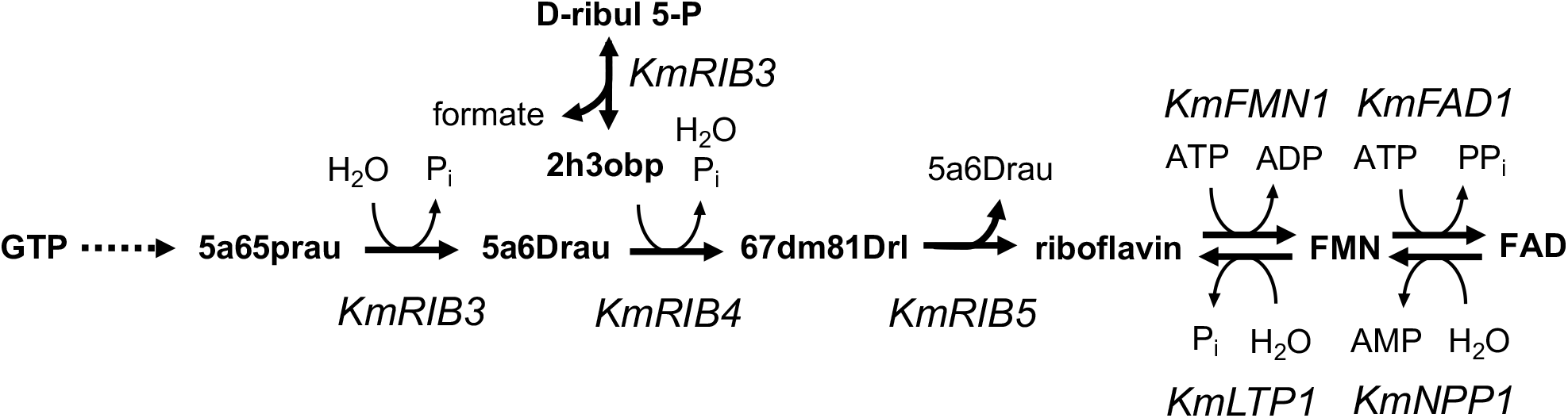
The riboflavin biosynthetic pathway. Abbreviations for metabolites: 2,5-diAm-4-hxy-6-(5-phosRsylAm)pymd (2,5-diamino-4-hydroxy-6-(5-phosphoribosylamino)pyrimidine), 2,5-diAm-6-(5-phos-D-rtylAm)pymd-4(3H)-one (2,5-diamino-6-(5-phospho-D-ribitylamino)pyrimidin-4(3H)-one), 5-am-6-(5-phosRtylAm)ucil (5-amino-6-(5-phosphoribitylamino)uracil), 5-am-6-(D-rtylAm)ucil (5-amino-6-(D-ribitylamino)uracil, D-ribul 5-P (D-ribulose 5-phosphate), 2h3obp (2-hydroxy-3-oxobutyl phosphate). Abbreviations for genes: KmRIB3 (3,4-dihydroxy-2-butanone-4-phosphate synthase), KmRIB4 (lumazine synthase), KmRIB5 (riboflavin synthase), KmFMN1 (riboflavin kinase), KmFAD1 (FAD synthetase), KmNPP1 (nucleotide pyrophosphatase/phosphodiesterase), KmLTP1 (putative protein phosphotyrosine phosphatase).

The lipid pool size remains the same at all three conditions. However, this assumption is based on arbitrary abundance value for myo-inositol (0.001 mmol/gDW/h), so one may find the different lipid pool sizes once the experimental values for myo-inositol are used. The excessive availability for other lipid components is suppressed by inactivation of the KmDGK1 gene, which is a known contributor to the increased heat sensitivity when overexpressed [30].

The significant decrease of flux found in deoxynucleotides was the shortage of dAMP. Due to inactivation of the *KmISN1* and *KmSDT1* genes, dAMP shortage cannot be compensated by salvaging other purine nucleotides. This suggest that the cell to conserve these building blocks. This coincides with the upregulation of genes responsible for DNA repair [31].

## Conclusions

We present iSM996, the first genome scale-metabolic model for thermotolerant yeast *K. marxianus*. This model is capable of predicting growth on the majority of known carbon and nitrogen sources and was proved to accurately predict the growth rates. In addition, the model contains several unique biosynthetic pathways for aroma compounds like 2-phenylethanol, phenethyl acetate or ethyl acetate. Finally, the iSM996 was used to construct three temperature specific models in YPD medium. The results suggested that at high temperatures the cell turn off more genes thereby introducing new auxotrophies and hereby utilizing as many resources as possible from the medium.Additional files

**Additional files**

Additional file 1: Model file in SBML format (XML 4727 kb)

Additional file 2: Model file in Excel format (XLSX 275 kb)

Additional file 3: Biomass composition details for overall components (Sheet 1), deoxyribonucleotides (Sheet 2), ribonucleotides (Sheet 3), amino acids (Sheet 4). Also included the model performance upon chemostat conditions for the growth rate (Sheet 5), oxygen consumption (Sheet 6) and carbon dioxide production (Sheet 7). References are included in the file. (XLSX 41 kb)

Additional file 4: List of inactive genes (Sheet 1) and corresponding reactions (Sheet 2) between 30D, 30DS, 30X and 45D conditions. (XLSX 16 kb)

Additional file 5: **Figure S1**. A radar chart showing the predicted potential for biomass precursors excessive production in 30D, 30DS and 45D conditions. As the magnitude is different for each metabolite, the relative production values are shown, where 100% indicates the largest production capacity between conditions. The data for the 30D condition is shown as the green polygon bordered with the dashed border while the corresponding data for the 30DS condition is in purple (dotted border) and the data for the 45D condition is in red (dot dash border) color. (a) Abbreviations: G6P (alpha-D-glucose 6-phosphate), F6P (beta-D-fructose 6-phosphate), E4P (D-erythrose 4-phosphate), R5P (D-ribose 5-phosphate), GAP (D-glyceraldehyde 3-phosphate), 3PG (3-phosphoglycerate), PEP (phosphoenolpyruvate), PYR (pyruvate), OXA (oxaloacetate), ACA (acetyl-CoA), 2OG (2-oxoglutarate), SCA (succinyl-CoA). (b) Abbreviations: AMP (adenosine monophosphate), CMP (cytidine monophosphate), GMP (guanosine monophosphate), UMP (uridine monophosphate), dAMP (deoxyadenosine monophosphate), dCMP (deoxycytidine monophosphate), dGMP (deoxyguanosine monophosphate), dTMP (thymidine monophosphate). (DOCX 1117 kb)

## Abbreviations

GEM: : Genome-scale metabolic model
FBA: : Flux balance analysis
YPD: : Yeast extract peptone glucose
YPX: : Yeast extract peptone xylose

## Acknowledgements

Not applicable

## Availability of data and materials

The iSM996 GEM together with the supporting information will be made publicly available on GitHub upon the acceptance of the paper.

## Authors’ contributions

SM, BJ and JN conceived the study. SM and BJ performed model reconstruction and validation. SM performed condition-specific model reconstructions and comparative analysis. SM, BJ and JN drafted, reviewed and approved the manuscript.

## Ethics approval and consent to participate

Not applicable

