## Additional File 5 for "Reconstruction and Analysis of a *Kluyveromyces marxianus* Genome-scale Metabolic Model"

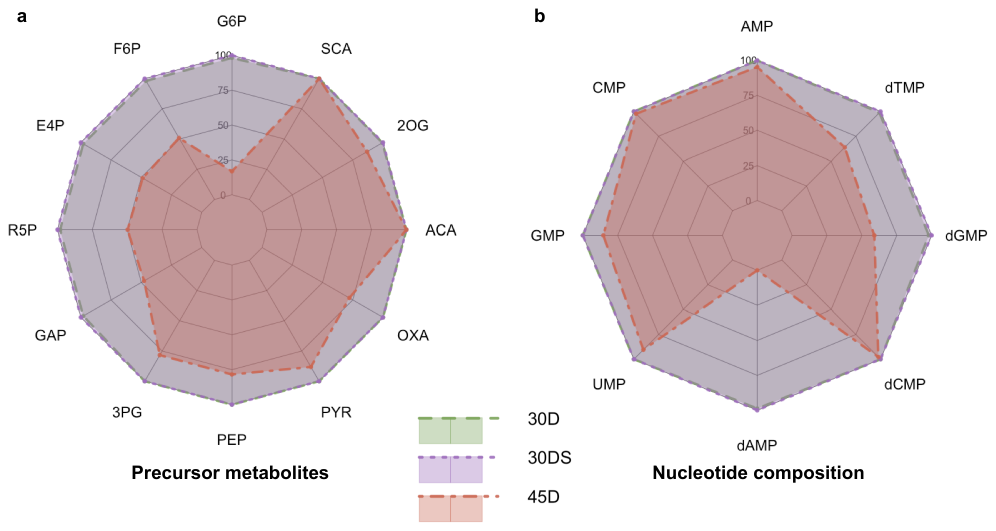
Figure S1. A radar chart showing the predicted potential for biomass precursors excessive production in 30D, 30DS and 45D conditions. As the magnitude is different for each metabolite, the relative production values are shown, where 100 % indicates the largest production capacity between conditions. The data for the 30D condition is shown as the green polygon bordered with the dashed border while the corresponding data for the 30DS condition is in purple (dotted border) and the data for the 45D condition is in red (dot dash border) color. (a) Abbreviations: G6P (alpha-D-glucose 6-phosphate), F6P (beta-D-fructose 6-phosphate), E4P (D-erythrose 4-phosphate), R5P (D-ribose 5-phosphate), GAP (D-glyceraldehyde 3-phosphate), 3PG (3-phosphoglycerate), PEP (phosphoenolpyruvate), PYR (pyruvate), OXA (oxaloacetate), ACA (acetyl-CoA), 2OG (2-oxoglutarate), SCA (succinyl-CoA). (b) Abbreviations: AMP (adenosine monophosphate), CMP (cytidine monophosphate), GMP (guanosine monophosphate), UMP (uridine monophosphate), dAMP (deoxyadenosine monophosphate), dCMP (deoxycytidine monophosphate), dGMP (deoxyguanosine monophosphate), dTMP (thymidine monophosphate).
